# An exploratory *in silico* study of the effect of a homogeneous static magnetic field on membrane dynamics and the CNGC6 ion channel of *Solanum lycopersicum* L

**DOI:** 10.64898/2026.06.06.730596

**Authors:** Camilo Tayac, J. Torres-Osorio, José Mauricio Rodas-Rodríguez

## Abstract

Magnetic treatment in tomato seeds (*Solanum lycopersicum* L.) has been studied as a biotechnological technique to induce a reduction in germination times and enhance plant development. However, the modified cellular mechanisms involved in the reduction of germination times or the improvement of development parameters are not yet clearly established. To explore a possible altered cellular mechanism, the effect of homogeneous static magnetic fields on the structure of the cyclic nucleotide-gated channel 6 (CNGC6), the modification in the organization of POPC lipids in the plasma membrane, and changes in calcium ion mobility were evaluated. For this purpose, coarse-grained molecular dynamics simulations were performed using the Martini 3 model in GROMACS, applying five different magnetic flux densities (0.000, 0.001, 0.010, 0.100, 1.000, and 10.000) T over 1 000 ns. The results showed an anisotropic effect in the longitudinal direction of the protein, which generated heterogeneous behavior among the chains of the homotetramer; this altered the conformation of the CNGC6 channel and modified the pore bottleneck. In contrast, no significant changes were observed in the conformational order of the POPC phospholipid chains. As a preliminary, single-replicate exploratory study, these results suggest that homogeneous static magnetic fields may induce specific structural modifications in the CNGC6 ion channel of *Solanum lycopersicum* L. without compromising the integrity of the lipid bilayer or the dynamics of ion transport within the analyzed timescale; these preliminary findings provide a molecular-level structural basis for future experimental and computational investigations of magnetic field effects on plant cyclic nucleotide-gated channels.

## 1. Introduction

Cyclic nucleotide-gated (CNGC) channels are specialized proteins found in various plant species and, in turn, in different plant and cellular structures [1]. These channels are relevant in the regulation and transport of Ca^2+^, and they acquire significance due to the role they play in intracellular signaling, the regulation of membrane potential, and multiple physiological processes. They are channels in which second messenger activation overlaps with voltage activation; they are located in the plasma membrane, chloroplast membrane, nuclear membrane, and vesicles [1]. There are reports proposing that static magnetic fields (SMFs) can modulate the structural and functional properties of the plasma membrane, including membrane potential, the gating dynamics of ion channels, and ATPase activity [2,3]. In this context, CNGC channels can be considered potential magnetosensors in plant cells under the influence of SMFs.

At a macroscopic level, magnetic seed treatment (MST) at 100 mT has shown positive effects on *Solanum lycopersicum* L., promoting germination and improving establishment and vegetative development [4]. Poinapen and his research team propose that the application of SMFs with intense magnetic flux densities (B) in tomato seeds can significantly increase the structural order of lipids at the cellular level, favoring a more rigid gel phase and reducing the proportion of lipids in the fluid phase of the cell membrane [5]. On the other hand, when applying an SMF to *Spinacia oleracea*, the Ca^2+^ flux across the plasma membrane was altered [6]. A physical mechanism that could explain the proposed effects at the cell membrane level is the torque generated by the interaction of cellular structures—such as lipid bilayers, microtubules, ɑ-helix proteins, and nucleic acids—with the magnetic field lines, which induces the spatial reorientation of the components, as described in [7] and specifically in vertebrates, the torque on the EGFR kinase monomer [8].

Despite the advances in the study of the exposure of plant systems to SMFs, a gap persists between the identification of the effects resulting from the application of MST and the understanding of the cellular mechanisms that are modified in the plasma membrane. In this context, CNGCs and, in particular, the CNGC6 of *Solanum lycopersicum* L., constitute an interesting structure for focusing *in silico* investigations due to their central role in transmembrane Ca^2+^ transport and in the regulation of multiple physiological processes. However, there are few studies on the possible structural alterations that these channels might undergo under the action of SMFs, nor how these modifications impact the interaction with membrane lipids and the associated ionic flux.

To contribute to the explanation of the effects that SMFs can generate on cell membrane components at a mesoscopic scale, it is necessary to delve deeper into whether the exposure of plant cellular structures to SMFs can lead to specific alterations. Therefore, this work studied the effect of exposure to a homogeneous SMF on: i) the structural behavior and pore geometry of the CNGC6 of *Solanum lycopersicum* L., ii) the behavior of POPC-type lipids in the plasma membrane, and iii) the behavior of Ca^2+^ and Cl^-^ion movement through the study system. To address this issue, coarse-grained (CG) molecular dynamics simulations were employed, seeking to combine computational efficiency and biological relevance.

## 2. Methodology

The process began with the construction of the study system, composed of a lipid bilayer, the CNGC6 channel from *Solanum lycopersicum* L. (henceforth referred to only as CNGC6), water beads, and Ca^2+^ and Cl^-^ion beads. Each system replica was subjected to the same protocol, which included three stages: (i) energy minimization, (ii) equilibration, and (iii) molecular dynamics production. The SMF was applied starting from the equilibration stage, maintaining the same steps on the system for the exposure simulations with five magnetic treatments. Subsequently, an analysis of the quantified data for each structure was performed to assess the generated effects.

### 2.1 Implementation of the Study System

A protein-membrane structure was configured, consisting of a CNGC6 channel embedded in a lipid bilayer, immersed in a Ca^2+^ and Cl^-^solution. The protein modeling was previously implemented by the authors of this work [9] and is available in ModelArchive under the code ma-pcoua. For the construction of the protein-membrane system, the orientation of the CNGC6 channel in the membrane was determined using the PPM 3.0 tool [10], which allowed for the proper positioning of the CNGC6 channel across the membrane. Based on this orientation, a lipid bilayer composed of a total of 2 683 POPC lipids was generated using the Martini Bilayer Maker tool [11,12], available in CHARMM-GUI [13], and applying the Martini 3 coarse-grained force field [14]. The system was solvated with 50 171 W-type water beads and 934 Ca^2+^ ions represented by SD-type beads; to maintain the electroneutrality of the solution, 1 912 Cl^-^ions were added. The implemented system is presented in Figure 1.

**Figure 1.**
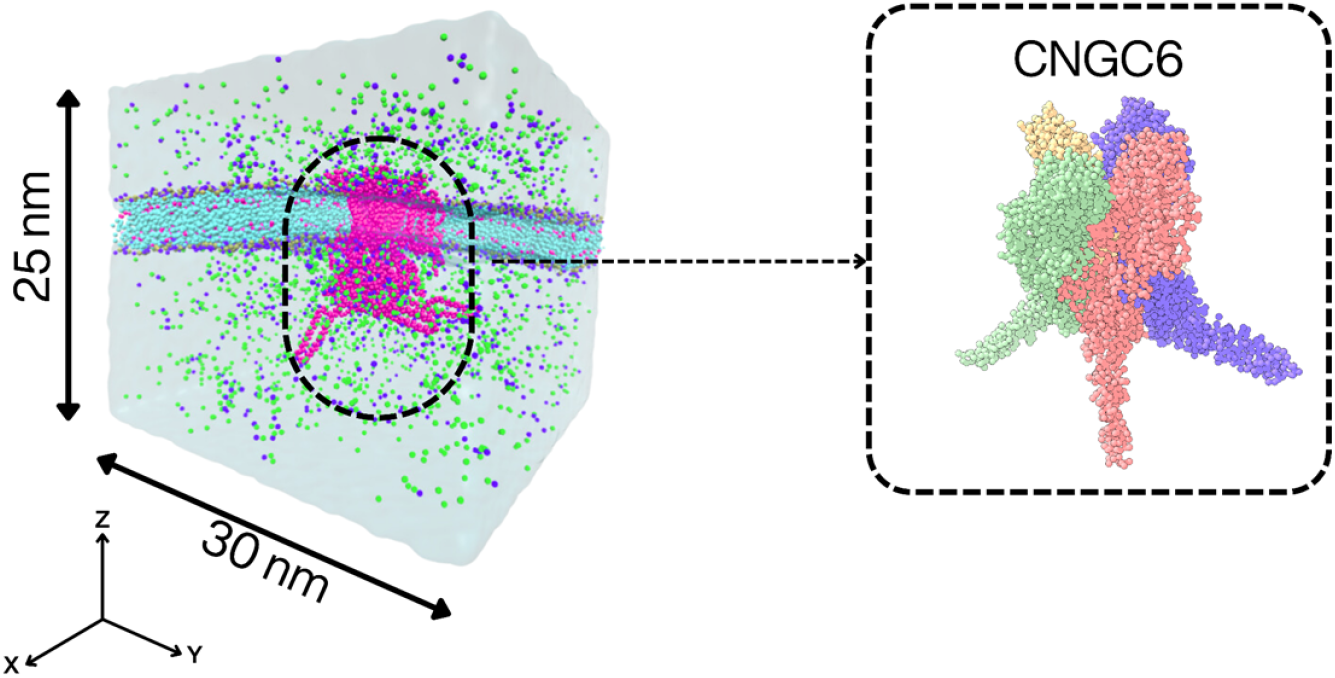
Representation of the CG system (∼ 205 kbeads); purple beads are Ca^2+^, green beads are Cl^-^, violet beads represent the CNGC6 channel, and aquamarine beads correspond to POPC lipids. In the protein inset, each color represents each of the four monomers of CNGC6.

Next, the entire system was subjected to two energy minimization stages, followed by five equilibration stages under NPT ensemble conditions (303.15 K and 1.0 bar), using the V-rescale thermostat and the C-rescale barostat. Subsequently, production molecular dynamics simulations were performed for 1 000 ns using GROMACS software [15], with the md-vv integrator, the V-rescale thermostat, and the Parrinello–Rahman barostat. During the production stage, system evolution information was stored every 100 integration steps. No position restraints were applied to any component of the system during the production runs, allowing the protein, lipids, and ions to evolve freely under the applied field conditions.

### 2.2. Molecular Dynamics Simulations under SMF Exposure

To evaluate the potential impact of the homogeneous SMF on the channel dynamics, a modified version of the GROMACS source code, developed by members of the same research groups involved in this work [16], was employed. Six independent simulations of 1000 ns each were performed: the first without a magnetic field (0.000 T) as a control, and five with SMF whose B values were selected following a base-10 logarithmic scale (0.001, 0.010, 0.100, 1.000, and 10.000) T. This methodological choice allowed for an analysis across four orders of magnitude to determine the system’s response across a range of B. All simulations were performed with the B vector in the Z-direction. Regarding statistical sampling, due to computational resource constraints, a single production run was performed per magnetic field condition; the observed differences between conditions should therefore be interpreted as preliminary exploratory trends, and the consistency of structural responses across all analyzed metrics supports the direction of the reported findings. Independent replicate simulations are recommended to establish full statistical significance in future work.

The duration of 1 000 ns for the production stage corresponds to criteria of statistical convergence and computational cost. In coarse-grained simulations with the Martini 3 force field, the effective temporal acceleration factor relative to atomistic models is estimated to be between 4X and 8X, which implies that 1 000 ns of CG simulation correspond, as a guideline, to an effective timescale on the order of microseconds. This interval allows for the sampling of multiple slow conformational transitions of the channel, including fluctuations of the CNBD domain and reorientations of transmembrane segments. Convergence was verified through the stabilization of the RMSD and radius of gyration profiles starting from 100 ns. Regarding the Parrinello–Rahman barostat: it was used exclusively during the production stage according to the recommendations of the CHARMM-GUI protocol for protein-membrane systems with Martini 3. Unlike the Berendsen barostat used in equilibration, which artificially damps pressure fluctuations, the Parrinello–Rahman barostat reproduces the correct pressure fluctuations of the statistical NPT ensemble and is the standard recommended by GROMACS documentation for production simulations with CG force fields [15].

### 2.3. Evaluation Parameters

To evaluate the effects corresponding to the five SMF values on the study system, molecular dynamics trajectories were analyzed using nine metrics and five computational tools from 100 ns onwards. For the CNGC6 channel, the structural behavior and pore geometry were analyzed. The positioning of POPC lipids and the orientational order of the lipid tails, corresponding to the sn1 and sn2 chains, were evaluated; regarding ions, the mobility of Ca^2+^ and Cl^-^was assessed. The trajectory preparation and structure selection protocols applied prior to these analyses are described below; the complete set of analyses, metrics, and tools employed is summarized in Table 1.

**Table 1:**
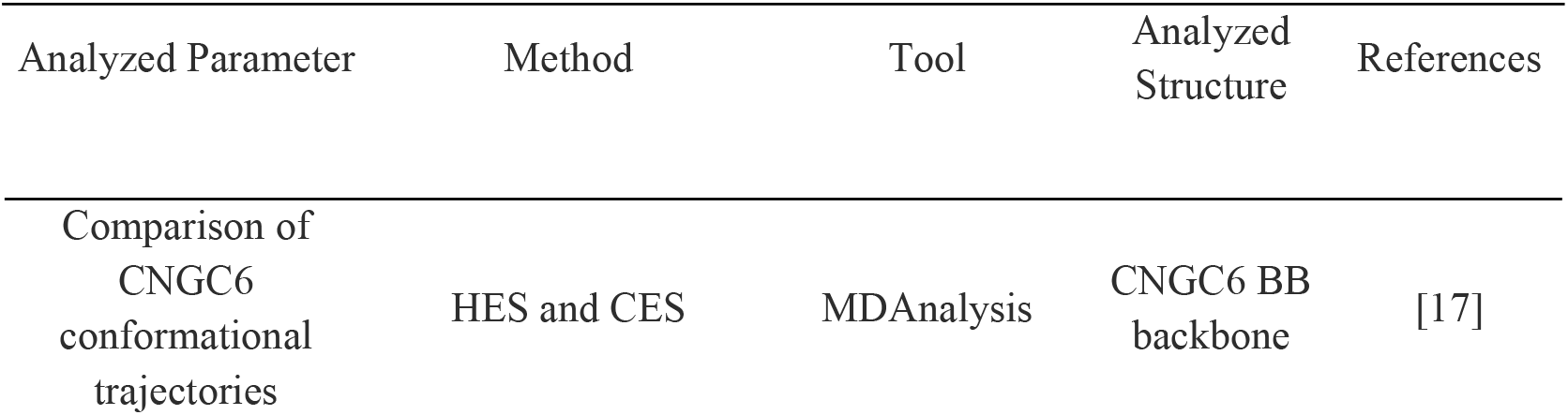
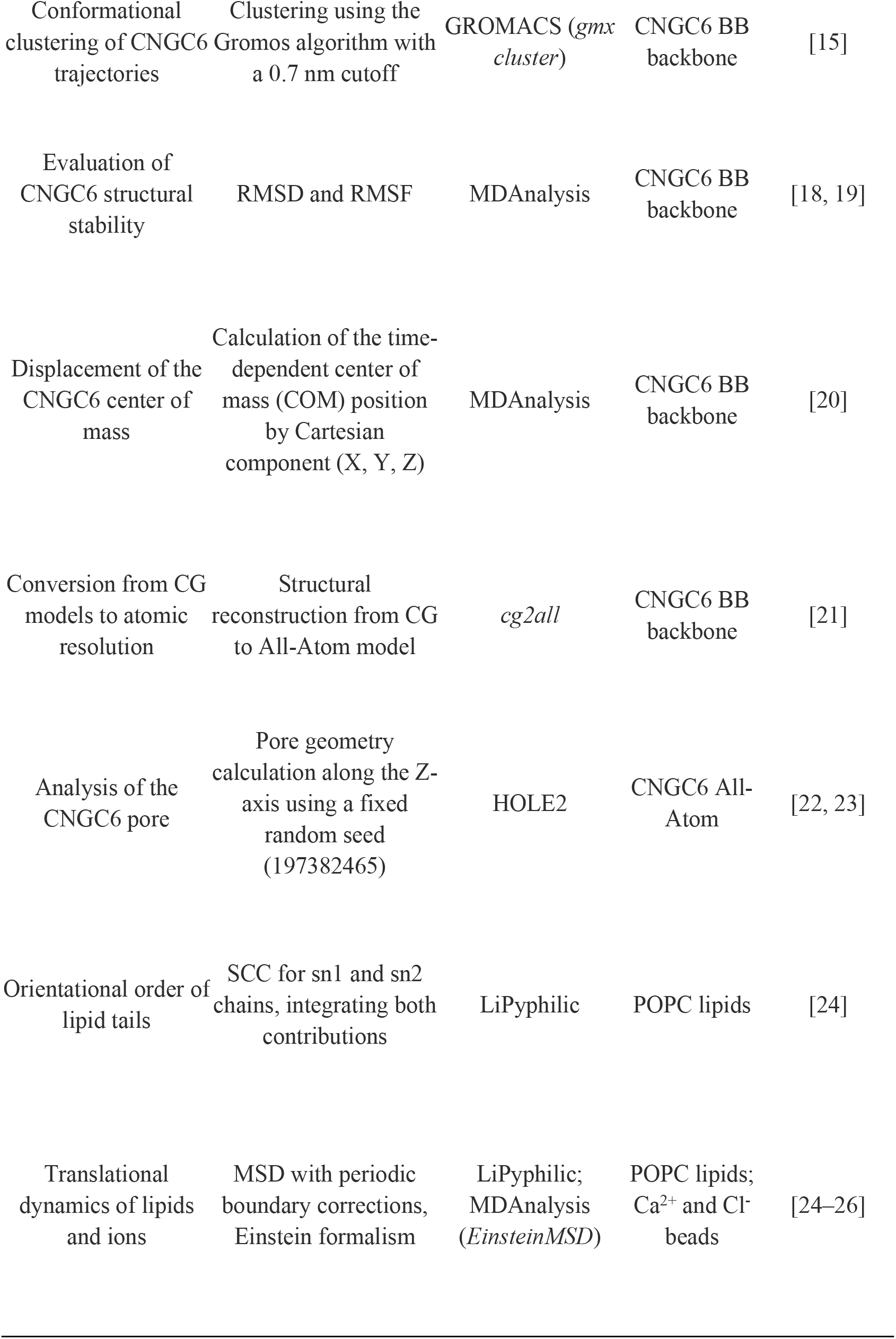
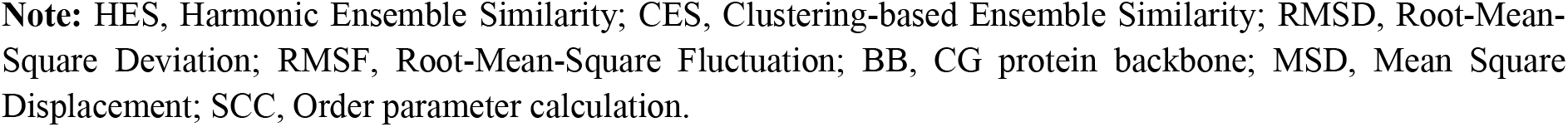
Metrics and tools used for the structural and molecular dynamics characterization of the study system exposed to a homogeneous SMF.

| Analyzed Parameter | Method | Tool | Analyzed Structure | References |
| --- | --- | --- | --- | --- |
| Comparison of CNGC6 conformational trajectories | HES and CES | MDAnalysis | CNGC6 BB backbone | [17] |
| Conformational clustering of CNGC6 trajectories | Clustering using the Gromos algorithm with a 0.7 nm cutoff | GROMACS (*gmx cluster*) | CNGC6 BB backbone | [15] |
| Evaluation of CNGC6 structural stability | RMSD and RMSF | MDAnalysis | CNGC6 BB backbone | [18, 19] |
| Displacement of the CNGC6 center of mass | Calculation of the time-dependent center of mass (COM) position by Cartesian component (X, Y, Z) | MDAnalysis | CNGC6 BB backbone | [20] |
| Conversion from CG models to atomic resolution | Structural reconstruction from CG to All-Atom model | *cg2all* | CNGC6 BB backbone | [21] |
| Analysis of the CNGC6 pore | Pore geometry calculation along the Z-axis using a fixed random seed (197382465) | HOLE2 | CNGC6 All-Atom | [22, 23] |
| Orientational order of lipid tails | SCC for sn1 and sn2 chains, integrating both contributions | LiPyphilic | POPC lipids | [24] |
| Translational dynamics of lipids and ions | MSD with periodic boundary corrections, Einstein formalism | LiPyphilic; MDAnalysis (*EinsteinMSD*) | POPC lipids; Ca<sup>2+</sup> and Cl<sup>-</sup> beads | [24–26] |

**Note:** HES, Harmonic Ensemble Similarity; CES, Clustering-based Ensemble Similarity; RMSD, Root-Mean-Square Deviation; RMSF, Root-Mean-Square Fluctuation; BB, CG protein backbone; MSD, Mean Square Displacement; SCC, Order parameter calculation.

Prior to analysis, trajectories were post-processed with GROMACS trjconv using `-pbc nojump -center -ur compact` to correct periodic boundary artifacts, followed by least-squares fitting of backbone (BB) beads to the reference structure (`-fit rot+trans`) using a group index generated with gmx make_ndx. The resulting fitted trajectories were then loaded into MDAnalysis for all subsequent analyses, with no additional coordinate corrections applied, as periodic boundary artifacts and translational drift had already been removed at the GROMACS level. For pore geometry analysis specifically, representative structures were obtained by clustering each production trajectory using `gmx cluster` with the linkage method and an RMSD cutoff of 0.2 nm; the centroid of the most populated cluster was selected for each condition, back-mapped to all-atom resolution using cg2all, and submitted directly to HOLE2 without further energy minimization to preserve the backbone geometry as determined by the CG simulation.

## 3. Results and Analysis

### 3.1. Comparison of conformational trajectories of CNGC6

Fig. 2 presents two complementary metrics for the comparison of conformational ensembles of the protein backbone. Panel (a) shows the Jensen–Shannon divergence (DJ-S) between pairs of trajectories; this metric, derived from information theory, quantifies the dissimilarity between probability distributions of the conformational states of the protein backbone, with values between 0 (total conformational identity) and 1 (maximum dissimilarity). Panel (b) presents the covariance matrices of the BB bead fluctuations between pairs of simulated conditions, in which the color code reflects the magnitude of the cross-covariances and identifies whether two trajectories share correlated movement patterns. Both tools are complementary: DJ-S evaluates the global overlap of the conformational space, while the covariance matrix allows for the identification of protein regions with differential correlations between conditions.

**Figure 2.**
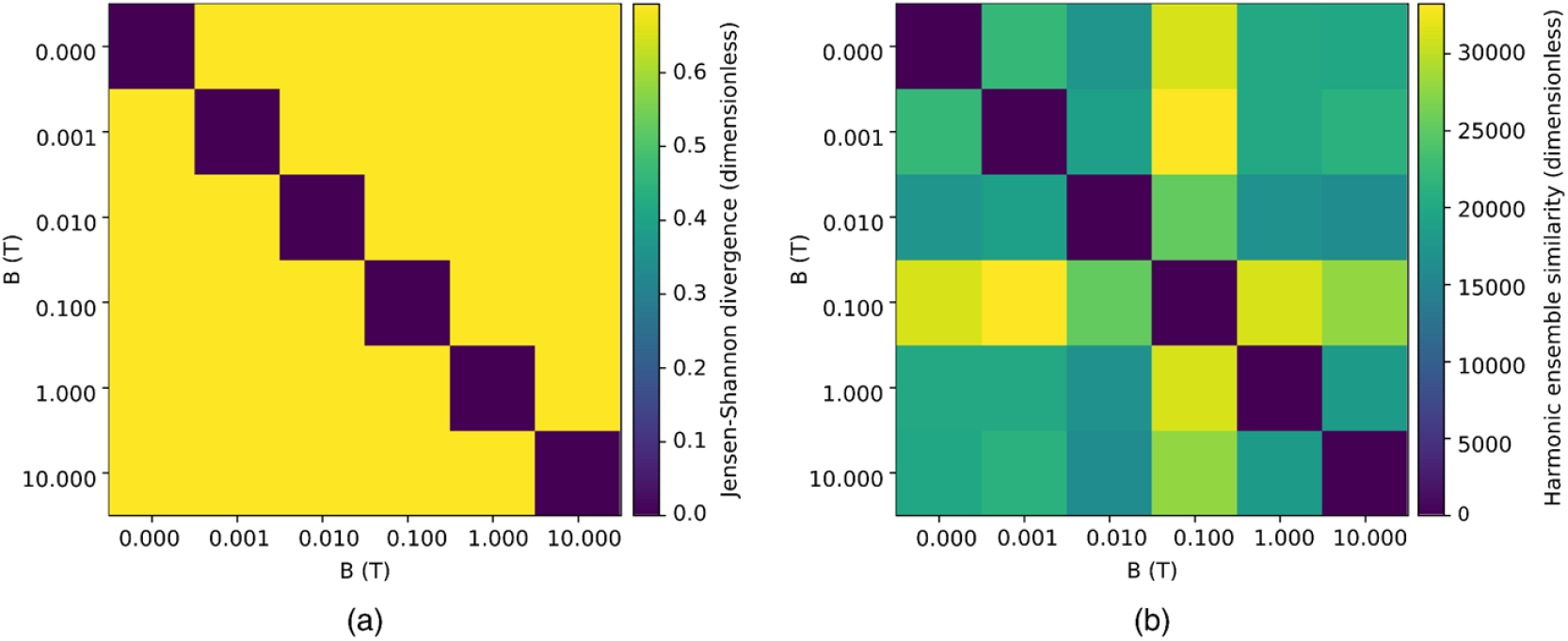
Comparison of the BB bead trajectories for five values of B using: (a) Jensen–Shannon divergence, (b) covariance matrices.

The trajectories of the backbone (BB) beads of CNGC6 as a function of B vary without a specific trend, as can be inferred from the analysis of Fig. 2. It is noteworthy that the Jensen– Shannon divergence between the conformational trajectories shows values greater than 0.6 in all comparisons (Fig. 2a), which indicates a low overlap between the most representative BB conformations for each value of B. This suggests that the interaction with the SMF introduces a bias in the conformational sampling of the system. While the stochastic fluctuations of the membrane continuously alter the position and the global tilt angle of the channel, each value of B favors a predominant orientation. The protein’s ɑ-helices experience a torque that seeks to align their longitudinal axes with the direction of the B vector, as previously proposed by some authors [7,8]. On the other hand, the covariance matrices of the BB fluctuations (Fig. 2b) present a heterogeneous pattern, although intermediate values (green) predominate, reflecting moderate differences in the trajectories. Specifically, exposure to 0.100 T exhibits a marked difference in the followed trajectory compared to the other conditions.

### 3.2. Structural Analysis of CNGC6

To analyze the behavior of the transmembrane region and the CNBD domain of the C-terminal region, which corresponds to residues 91 to 610 of each channel monomer, the dynamics within the channel were evaluated based on the BB positions, the variation of BB positions among the chains, and the compaction of the homotetramer.

By evaluating how the BB positions change in each replica subjected to SMF using Fig. 3, it is observed that for all values of B, the RMSD of the BBs in this region is higher than that recorded for the control, indicating that the SMF can modify the selected structure. Specifically, the protein subjected to exposure at 0.100 T presented the greatest dispersion in RMSD values, although the median matches that of 1.000 T. On the other hand, the protein exposed to 0.001 T showed the highest RMSD, while the lowest median was observed with the 0.010 T exposure.

**Figure 3.**
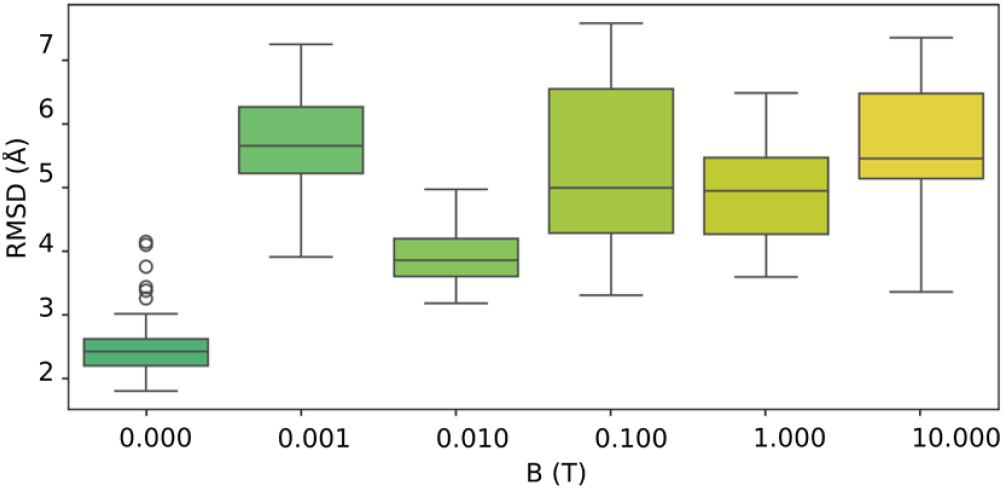
RMSD analysis of the BB beads for residues 91 to 610 of CNGC6.

Now, when analyzing the variation in the position of the BBs among the CNGC6 chains, it is noteworthy that there is also an affectation under SMF exposure for this parameter. Comparing the RMSF per chain (Fig. 4), it is observed that there are no variations between chains at exposures of 1.000 T and 10.000 T, as the means are similar. However, at 0.001 T, 0.010 T, and 0.100 T, variations in the median RMSF are detected, particularly in chain C, and in the case of 0.100 T, also in chain D. The RMSF per chain does not describe a monotonic relationship with B; it is a non-linear response reflecting a selective sensitivity dependent on the field value. At 1.000 T and 10.000 T, the response per chain is homogeneous, with similar means among chains A, B, C, and D, which could be misinterpreted as an absence of effect. However, this homogeneity does not imply an absence of perturbation, but rather the presence of a uniform magnetic confinement regime acting symmetrically on all monomers. At fields from 0.001 T to 0.100 T, the differential diamagnetic anisotropy between monomers, modulated by stochastic membrane tilt fluctuations, generates torques of different magnitudes on each chain, producing the observed intratetrameric heterogeneity. The apparent equivalence of the chains at high fields, therefore, constitutes a magnetic saturation regime rather than an absence of interaction.

**Figure 4.**
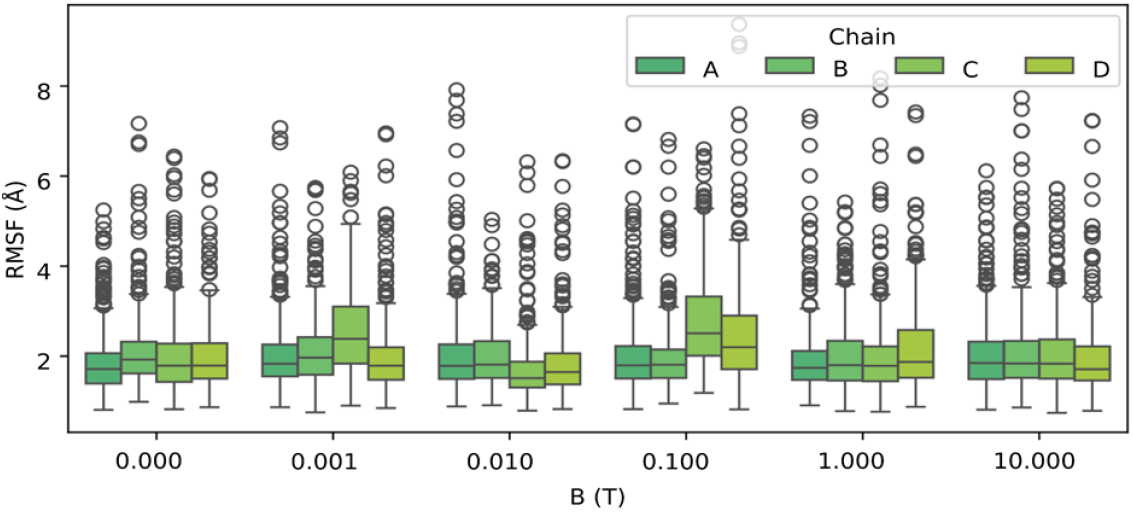
RMSF analysis of the beads corresponding to residues 91 to 610 of CNGC6 for each chain of the homotetramer. For each value of B, the response corresponding to each monomer is represented.

Regarding the degree of structural compaction of CNGC6, this factor is also affected by specific B values depending on the direction considered. For the R_g_ along the X-axis (Fig. 5a), no changes are detected except for an increase in the parameter under 0.100 T exposure, indicating that the protein is more expanded compared to the control in this direction. In contrast, the R_g_ along the Y and Z axes (Fig. 5b, c) show different behaviors; regarding the Y-axis, the structure appears more compact for all analyzed B values. Along the Z-axis, greater compaction is observed under 0.100 T, 1.000 T, and 10.000 T. When evaluating the overall R_g_ for each B value (Fig. 5d) compared to the control system, it is evident that the structure adopts a more expanded conformation under B values of 0.001 T and 0.100 T, defined by an increase in the R_g_. Conversely, under exposures of 0.010 T, 1.000 T, and 10.000 T, the protein shows a higher degree of compaction relative to the condition without SMF. This global expansion specifically observed at 0.001 T and 0.100 T is directly linked to the dynamic asymmetry described in the previous paragraph: the increase in the flexibility of chains C and D prevents compact packing of the tetramer.

**Figure 5.**
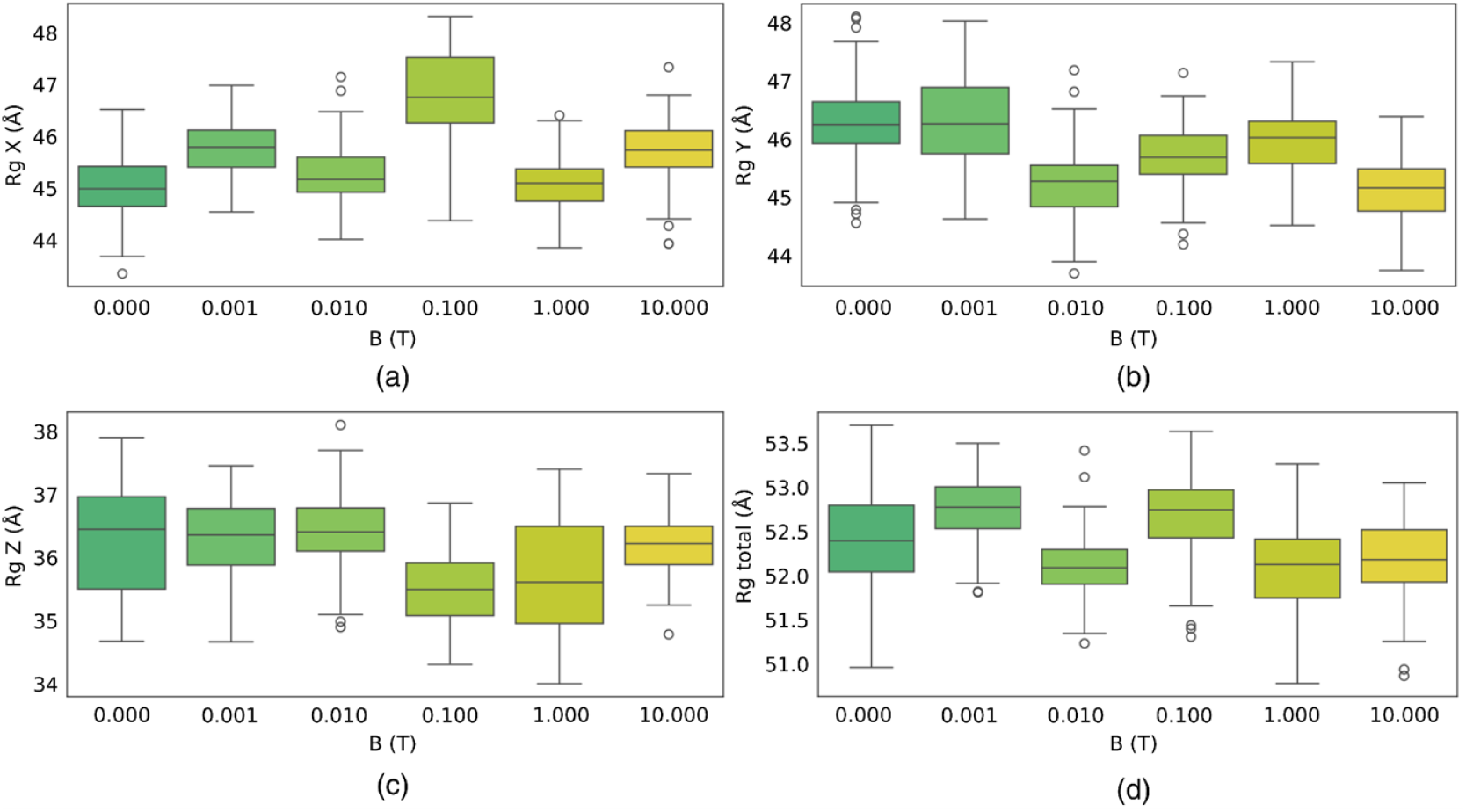
Radius of gyration (R_g_) analysis of the CNGC6 BB residues exposed to five values of B. (a) X-component. (b) Y-component. (c) Z-component. (d) Overall R_g_.

Since a general modification of the quaternary structure was determined, we proceeded to evaluate whether there was an effect at the level of specific chemical groups by analyzing the position of individual residues. The analysis of the average RMSF median per residue type (non-polar: Gly, Ala, Val, Leu, Ile, Pro, Cys, Met, Phe, Trp; polar: Ser, Thr, Asn, Gln, Tyr; acidic: Asp, Glu; basic: Arg, Lys, His) revealed no significant differences between SMF exposure conditions. These results suggest that, while magnetic exposure induces modifications in the global topology of the CNGC6 channel, such an effect does not derive from a selective interaction based on the chemical nature of the amino acids, but rather from the mechanical response of the protein backbone. Consequently, this figure has been removed from the main body of the manuscript, and the results are described in this paragraph to reduce the total number of figures in accordance with the reviewer’s recommendation.

When evaluating the center of mass (COM) of the BBs composing the channel in the X and Y directions (Fig. 6a), no variation in the median is observed compared to the control — an expected outcome of the rot+trans fitting applied during trajectory post-processing, which removes global translational drift in the transverse plane by construction. In contrast, a displacement is detected along the Z-axis; while in the absence of SMF, the center of mass is located around 140 Å measured from the base of the intracellular system, under the influence of SMFs it shifts by 35 Å toward the intracellular region, reflecting genuine vertical repositioning of the channel consistent with the membrane constraining lateral motion while permitting axial displacement. Analyzing this effect per chain reveals that the displacement is not homogeneous, indicating that despite the channel’s structural symmetry, the response to the SMF varies among chains. Exposures to 0.100 T and 1.000 T show a generalized variation of the center of mass across all chains (Fig. 6b). At 0.100 T, the chain with the greatest displacement is chain B, whereas at 1.000 T, the displacement occurs in chain D. In contrast, at exposures of 0.001 T and 10.000 T, a change in the COM median is observed only in chain A, while chains B, C, and D maintain similar values to each other.

**Figure 6.**
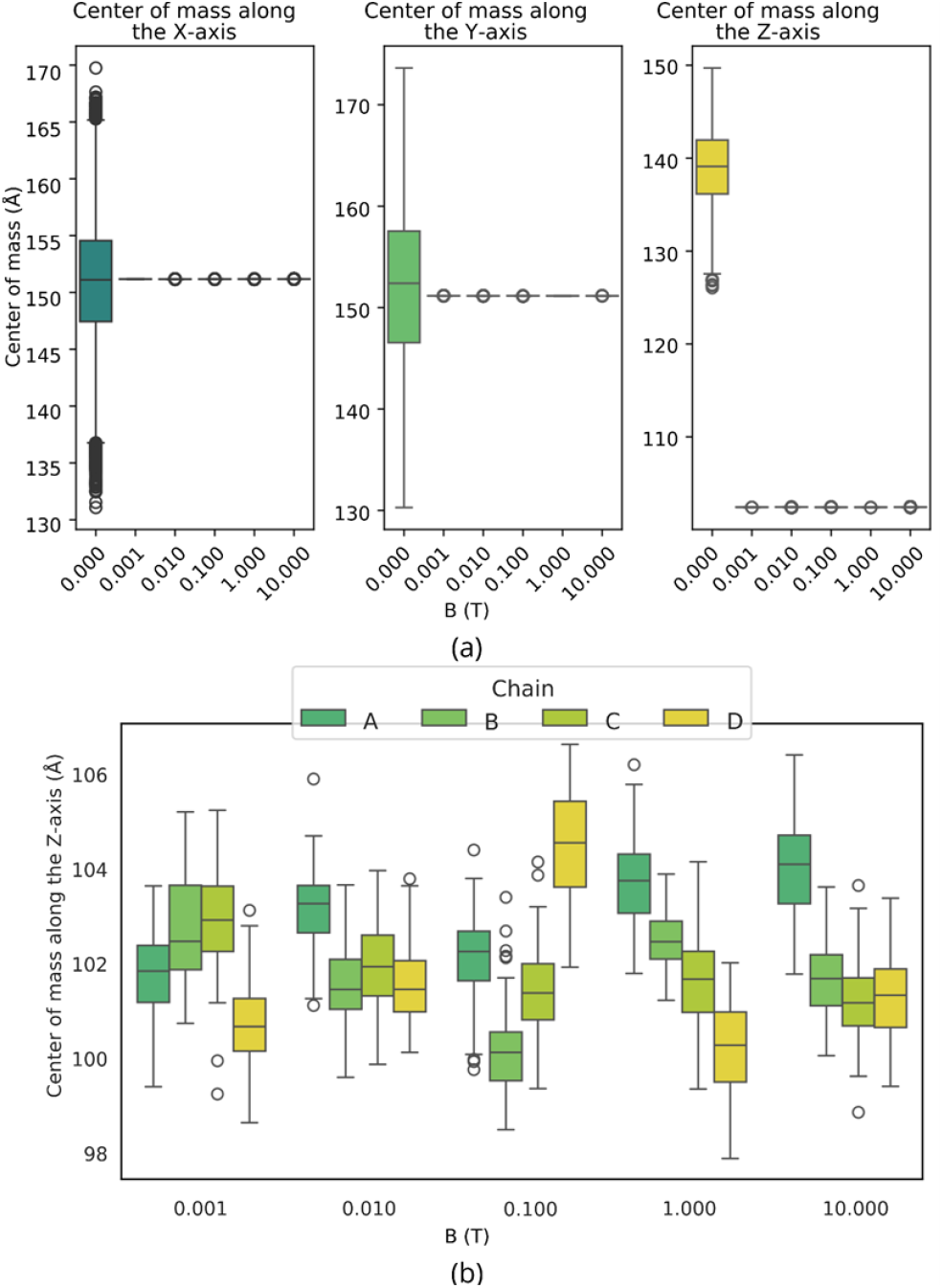
Center of mass (COM) variation of the CNGC6 channel under different B values. (a) Center of mass variation of the channel along the X, Y, and Z axes. (b) Center of mass variation along the Z-axis for each channel chain.

The integral analysis of conformational stability suggests that the interaction with the SMF does not constitute a uniform perturbation, but rather induces a complex structural reorganization governed by the anisotropic response of the system. Although the generalized increase in RMSD values evidences a global deviation relative to the control, the detailed analysis of the RMSF reveals an intrinsically heterogeneous response, which mostly affects chains C and D of the homotetramer, especially under exposures of 0.001 T and 0.100 T. This internal asymmetry translates into a mesoscopic deformation whose strongest evidence is manifested in the center of mass behavior.

The analysis demonstrates that the interaction with the SMF is restrictive and markedly directed along the Z-axis. While the system preserves its positional stability in the transverse plane, defined by the X and Y axes, it is fundamental to highlight that the SMF not only induces a protein displacement but also substantially modifies the vertical fluctuation dynamics. In the condition without SMF, the center of mass exhibits a wide dispersion, indicating high freedom of oscillation along the Z-axis. In contrast, under SMF, the variability is reduced, resulting in compact distributions. This behavior is consistent with the presence of a magnetic confinement or anchoring effect that limits the natural fluctuations of the channel in the direction of the B vector.

This decoupling, combined with the restriction of vertical mobility, shows that the system undergoes a differential alteration of its dynamics, where magnetic forces suppress longitudinal oscillations while preserving the spatial arrangement in the transverse plane. The convergence of these results—including heterogeneity among monomers, confinement along the Z-axis, and anisotropic deformation—indicates that the underlying physical mechanism corresponds to a vectorial interaction dependent on the spatial orientation of the magnetic field. Since the intrinsic stochastic fluctuations of the membrane induce dynamic variations in the protein tilt, an asymmetric angular deviation occurs between monomers, leading to the action of torques of differential magnitude on each monomer. As a consequence of this effect, a heterogeneous structural reorganization emerges. Given that this observed phenomenon could be a consequence of the B vector direction, it is suggested for future research to apply B with different vectorial directions, which would allow for evidence of whether the anchoring persists or is modulated under different conditions.

### 3.3 Pore Geometry Analysis

The analysis of the pore radius of the CNGC6 channel along the ion transport axis (Z-axis), under different B values, indicated a shift in the bottleneck position toward the extracellular region. This effect was observed in the structure at an All-Atom resolution, obtained from the conversion of the most representative BB position identified through cluster analysis for each simulated condition. The position of the pore bottleneck radius under 0.001 T, 1.000 T, and 10.000 T is similar to the control; however, with exposure values of 0.010 T and 0.100 T, a shift in the pore bottleneck position toward the extracellular space is recorded (Fig. 7), suggesting the relocation of the BBs representing the pore-forming residues. Furthermore, under exposures to 0.100 T and 1.000 T, pore radius peaks smaller than the Ca^2+^ radius are identified, indicating a possible restriction of ion flux across the membrane. In contrast, the protein exposed to 10.000 T presents a bottleneck with a radius larger than that of Ca^2+^, suggesting that this B value would not limit ion passage. These results suggest that intermediate B values of 0.010 T and 0.100 T could induce variations in the pore radius of the CNGC6 channel, providing a structural basis for a mechanical hypothesis that requires future experimental validation. Although back-mapped structures were used without further energy minimization — preserving backbone geometry as determined by the CG simulation — absolute pore radius values should be interpreted with caution; the positional shift of the bottleneck along the Z-axis, governed primarily by backbone geometry, is nonetheless considered a reliable indicator of SMF-induced structural rearrangement.

**Figure 7.**
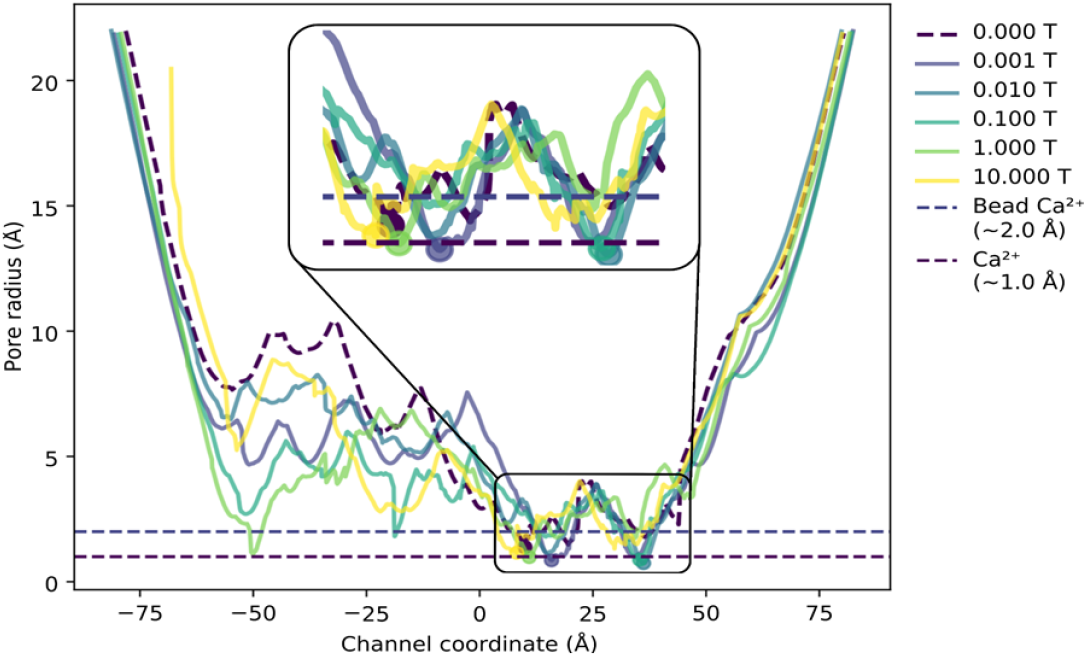
Pore radius profile along the Z-axis for the most representative structures of the CNGC6 channel under different B values. The blue dashed line indicates the radius of the Ca^2+^-type bead (Martini 3), and the purple dashed line indicates the Ca^2+^ radius, used as references to evaluate the potential pore permeability to Ca^2+^.

### 3.4. Analysis of POPC lipid and Ca^2+^ beads

SMF does not affect the reorganization of lipid tails or the mean movement of lipids in the bilayer. The ordering of the beads composing the sn1 and sn2 lipid tails of POPC during the 1000 ns simulation corresponds to a fluid state in all SMF exposure cases, with chain order parameter (SCC) values remaining between 0.30 and 0.34, with no significant differences between the different applied magnetic flux densities. Complementarily, the analysis of lipid bead movement through Mean Square Displacement (MSD) also showed no substantial differences among the studied B values, with the exception of the 0.001 T exposure, in which the MSD curve presents a transient elevation during the first few hundred nanoseconds before stabilizing into a regular diffusive behavior. These results can be attributed to the fact that the lipid tails, due to their structural orientation in the membrane, are partially aligned with the direction of the B vector applied in the Z-direction; however, this alignment is not sufficient to induce changes in the conformational order or the translational dynamics of the lipids at the evaluated timescales.

Regarding the movement of Ca^2+^ and Cl^-^ions, the average MSD analysis indicates that SMF does not induce substantial changes in their global translational mobility across the simulated trajectories in any of the evaluated magnetic exposure conditions. For both ionic species, the MSD curves, which represent the temporal evolution of the average displacement of the ions, show a progressive increase over time, characteristic of diffusive behavior in aqueous solution, with no evidence of saturation or spatial confinement that could be attributed to the SMF. For both ionic species, the 0.001 T exposure presents dynamics very similar to those observed in the control condition without a magnetic field, while the other magnetic flux densities tend to show slightly lower MSD values, although these differences do not reach a magnitude sufficient to attribute a direct and sustained effect of SMF on ionic diffusion in solution.

It is important to specify that the translational diffusion coefficient obtained through MSD exclusively characterizes ionic dynamics in solution and is conceptually distinct from the transmembrane ionic conductance of the channel. Estimating conductance through the pore would require calculating the Potential of Mean Force (PMF) along the pore axis, using techniques such as steered molecular dynamics (steered MD) or metadynamics, in combination with explicit transmembrane voltage conditions; all these methodologies are outside the scope of the present experimental design. Therefore, the absence of significant changes in ionic MSD confirms only that SMF does not alter the free and non-specific diffusion of ions in solution under the simulated conditions; the possible modulation of transmembrane ionic flux through the CNGC6 pore constitutes a hypothesis that depends solely on the documented pore geometric changes in section 3.3 (bottleneck displacement and effective radius variation), and not on alterations in free ionic mobility. Together, these results indicate that the interaction mechanism of SMF with the protein-membrane-ion system acts selectively and specifically on the macromolecular protein structure, leaving the diffusive dynamics of the ions dissolved in the aqueous medium invariant under the evaluated magnetic exposure conditions.

### 3.5 Analysis of responses relative to B values

The analysis of the evaluated metrics presents differentiated behaviors regarding the 0.001 T and 0.100 T values. The CNGC6 channel experiences conformational alterations characterized by an increase in R_g_, heterogeneity in the fluctuations of the backbone (BB) bead positions among the monomers, and, specifically at 0.100 T, a shift of the pore bottleneck toward the extracellular space accompanied by a decrease in the effective pore diameter, pointing to a possible restriction for Ca^2+^ transport. These conformational changes align with theoretical models in which diamagnetic anisotropy generates mechanical tension forces capable of modifying the open probability and pore geometry [27]. This indicates that the protein undergoes a destabilization of its quaternary symmetry and a loss of the base structural rigidity induced by the SMF.

In contrast, exposures to 0.010 T, 1.000 T, and 10.000 T induce a compaction of the global structure. Nonetheless, even under these higher compression conditions, an anisotropic response persists, evidenced by the systematic displacement of the center of mass along the Z-axis. This response supports the hypothesis proposing that, at specific B values, the magnetic interaction energy is sufficient to compete with the internal forces stabilizing the macromolecule’s quaternary structure [28]; however, the extrapolation of these structural effects to functional modifications in living cellular systems requires independent experimental validation, as the present study does not include transmembrane voltage conditions, intracellular regulatory factors, or the complexity of the actual cellular environment.

The overall analysis indicates that the CNGC6 channel exposed to 0.100 T presents a response derived from its magnetic anisotropy. This B value not only induces the clearest perturbations in the homotetramer symmetry—impacting chains C and D—but also constitutes a condition capable of simultaneously generating a global expansion of the structure (evidenced by the increase in R_g_), a bottleneck shift toward the extracellular space, and a restriction associated with the effective pore diameter. Consequently, exposure to 0.100 T can be assumed as the value at which the magnetic interaction overcomes the channel’s base structural stability, suggesting that at this level of magnetic exposure, the greatest potential for functionally modulating CNGC6 conductance would occur. This response at 0.100 T could be the basis for the cellular modification related to macroscopic behaviors that have reported physiological effects in tomatoes at 0.100 T [4, 29, 30].

### 3.6. Study Limitations

The present study has four principal limitations that should be considered when interpreting the results. First, due to computational resource constraints, only a single production run was performed per magnetic field condition; the observed inter-condition differences cannot be unambiguously attributed to the applied SMF rather than stochastic sampling variability, and independent replicate simulations are required to establish statistical significance. Second, the coarse-grained Martini 3 force field represents atoms as isotropic beads, precluding explicit representation of electronic anisotropy tensors required for a rigorous treatment of diamagnetic anisotropy; the structural changes observed are therefore best interpreted as emergent collective responses of the model. Third, all simulations were performed in the absence of a transmembrane electric field; the physiological relevance of the reported conformational changes remains to be established under voltage-clamp conditions representative of plant cell membranes. Fourth, pore geometry analysis was derived from back-mapped all-atom structures without further energy minimization, to preserve the backbone geometry determined by the CG simulation; absolute pore radius values should therefore be interpreted with caution, whereas the positional shift of the bottleneck along the Z-axis — determined primarily by backbone geometry — is considered a more reliable indicator of SMF-induced structural rearrangement.

## 4. Conclusions

Homogeneous SMFs appear to induce specific, B-value-dependent structural modifications on the CNGC6 ion channel of *Solanum lycopersicum* L. within the simulated timescales. These effects are evidenced in the positions of the BB beads, the pore geometry, and the bottleneck radius, without altering the organization of the POPC lipid bilayer or the ionic diffusion in solution. The observed conformational changes could be related to impairments in transmembrane ionic diffusion due to modifications in the channel’s pore bottleneck radius; this hypothesis requires validation through specific simulations with explicit transmembrane voltage or Potential of Mean Force (PMF) calculations.

Molecular dynamics simulations with different B values reveal varying behavior in the channel conformation, particularly along the Z-axis. These effects are dependent on B values, specifically 0.100 T, which perturbs chains C and D of the homotetramer. Furthermore, a potential impact on Ca^2+^ permeability is evidenced, associated with the displacement of the ion pore bottleneck under 0.010 T and 0.100 T. No substantial changes were observed in lipid organization or the mobility of lipids and ions, suggesting that the homogeneous SMF acts primarily on the protein through effects entailing a response associated with diamagnetic anisotropy. These findings constitute a hypothetical basis for future research to experimentally evaluate the functional modulation of CNGC6 conductance by SMF in complete biological systems, including validation in models with higher structural resolution and more representative physiological conditions. Given the exploratory and single-replicate nature of this study, all reported trends should be regarded as preliminary structural hypotheses requiring confirmation through independent replicate simulations and experimental validation.

## Data Availability Statement

All simulation input files, GROMACS parameter files (.mdp), force field topology files (.top, .itp), initial system coordinates (.gro), and Python analysis scripts used in this study will be deposited in a public repository (Zenodo) upon acceptance of the manuscript and will be freely available. Raw molecular dynamics trajectory files (.xtc) are available from the corresponding author upon reasonable request due to their large size.

## CRediT authorship contribution statement

**Camilo Tayac** obtained and compiled all the data used in the study, and also participated in the selection of reference articles, information review, drafting of the original manuscript, final structuring of the document, and organization of the content. **J. Torres-Osorio** contributed to data review and interpretation, drafting of the original manuscript, final structuring of the document, organization of the information, and data verification. **José Mauricio Rodas-Rodríguez** participated in data review and interpretation, information organization, and manuscript revision.

## Funding

No funding was received.

## Data availability

The data are uploaded to GitHub

## Declaration of generative AI and AI-assisted technologies in the writing process

During the preparation of this work the authors used Gemini in order to improve the grammar and language of the manuscript to enhance overall readability.

## Notes

### Competing Interest Statement

The authors have declared no competing interest.

